# Emergence of Bow-tie Architecture in Evolving Feedforward Networks

**DOI:** 10.1101/2023.03.28.534501

**Authors:** Thoma Itoh, Yohei Kondo, Kazuhiro Aoki, Nen Saito

## Abstract

Bow-tie architecture is a layered network structure that has a narrow middle layer with multiple inputs and outputs. Such structures are widely seen in the molecular networks in cells, suggesting that a universal evolutionary mechanism underlies the emergence of bow-tie architecture. The previous theoretical studies have implemented evolutionary simulations of the feedforward network to satisfy a given input–output goal, and proposed that the bow-tie architecture emerges when the ideal input– output relation is given as a rank-deficient matrix with mutations in network link strength in a multiplicative manner. Here, we report that the bow-tie network inevitably appears when the link intensities representing molecular interactions are small at the initial condition of the evolutionary simulation, regardless of the rank of the goal matrix. Our dynamical system analysis clarifies the mechanisms underlying the emergence of the bow-tie structure. Further, we demonstrate that the increase in the input–output matrix facilitates the emergence of bow-tie architecture even when starting from strong network links. Our data suggest that bow-tie architecture emerges as a side effect of evolution rather than as a result of evolutionary adaptation.

**Author Summary:** Many biological networks including gene regulatory networks, metabolic networks, and signaling networks, show a characteristic hierarchical structure known as the bow-tie architecture. This architecture consists of a narrow middle layer with multiple inputs and outputs. Understanding why bow-tie architecture has universally appeared through evolution may provide insight into the design principle of a network within a cell. The universality of the bow-tie structure has so far been explained by the adaptive advantages such as high evolvability and capability to classify the inputs. However, our computer simulation demonstrates that the bow-tie structure inevitably emerges even without functional advantages when the molecular interactions within the network are weak at the initial condition of the evolution. We also demonstrate that an increase in the number of inputs (i.e., receptors) and outputs (i.e., downstream genes) leads to the emergence of bow-tie architecture, even when evolution starts from a strong molecular interaction. Although many previous studies have discussed the adaptive properties of bow-tie architecture, our findings suggest that bow-tie architecture is a byproduct of evolution rather than a result of evolutionary adaptation.

## Introduction

Many metabolic, signaling, and gene regulatory networks have hitherto been reported to exhibit bow-tie architecture, also known as an hourglass structure system [1], which is characterized by hierarchical network structures with a narrow middle layer, multiple inputs and outputs. Bow-tie architecture receives a variety of inputs, and then converges signals into a few middle-layer components known as waists, cores, or knots [2]. Subsequently, the waist components regulate a wide range of downstream outputs [1] [3]. Bow-tie architecture has been found in metabolic networks [4] [5], gene regulatory networks [6] [7] [8] [9], signaling networks [10] [11] [12] [13] [14] [15] [16], and the neural network of the human visual system [2] (Fig 1). In the metabolic networks, various nutrients are decomposed into 12 universal precursors including G6P, F6P, PEP, PYR, AKG, and ACCOA, [4] [5], which are then used to generate abundant biomasses (e.g., proteins, fatty acid, and carbohydrates). The gene regulatory network involved in the developmental programs of *Drosophila* also contains bow-tie architecture; the multiple patterning genes, such as *Hox*, *wingless*, *EGF-R*, *hedgehog*, and *Notch*, induce the expression of a few transcription factors, such as *shavenbaby* and *scute*, which then regulate a myriad of downstream genes [6]. Bow-tie architecture is also found in intracellular signaling networks. For example, vast number of G-protein coupled receptors (GPCRs) encoded in the human genome primarily activate only a handful of G proteins followed by the regulation of the calcium ion or cAMP concentration, which nonetheless lead to the induction of diverse cellular phenotypes [17] [18] [19]. Bow-tie architecture has also been reported in non-biological networks, such as internet protocols [20] [21], railroad transportation systems [3], and so on. This universality of the bow-tie architecture implies the existence of the common design principles underlying these systems, and thus it is important to elucidate how and why the bow-tie architecture emerged.

**Fig 1.**
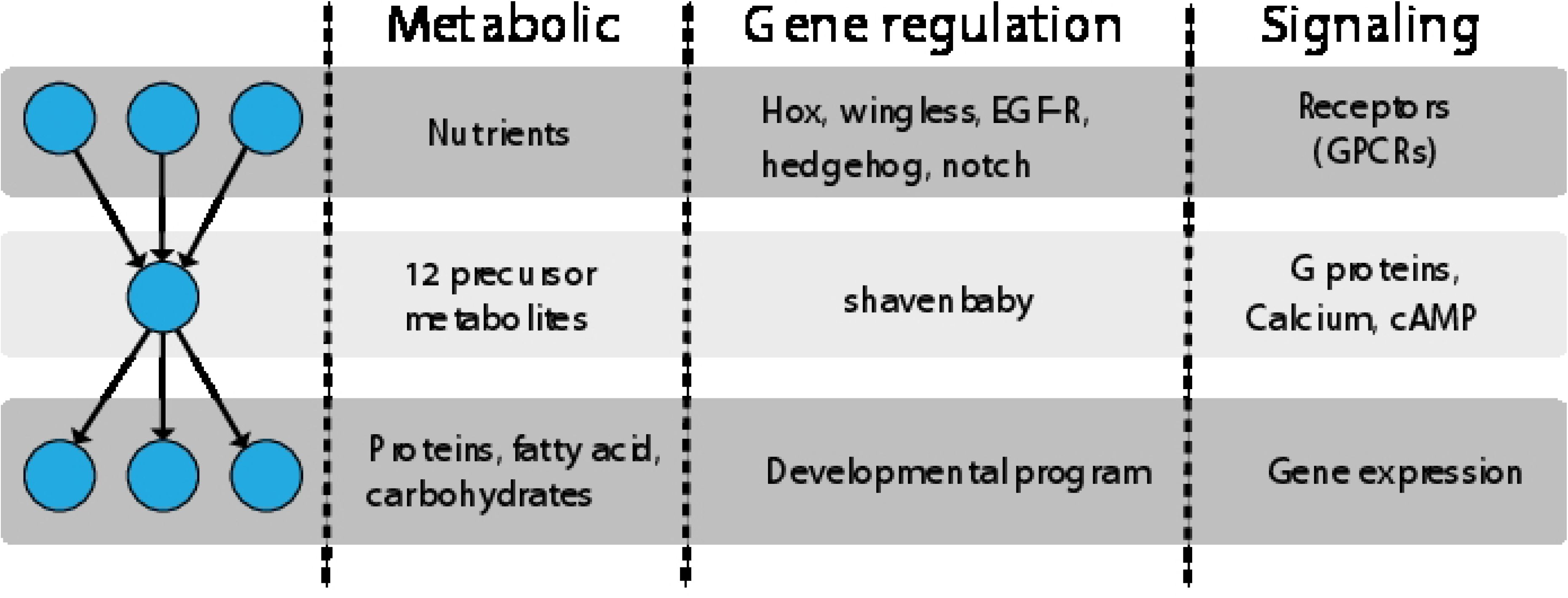
Universality of bow-tie architecture in a biological network. Bow-tie architecture is a hierarchical network having a narrow middle layer. A schematic of the typical bow-tie architecture is shown in the far left column. The circles (nodes) represent molecules and the arrows represent reactions or information flows. The other columns provide examples of bow-tie architecture in a metabolic network, gene regulatory network, and signaling network.

Despite the ubiquity of bow-tie architecture in living systems, the driving forces behind its emergence are still being debated. Kitano et al. have argued that bow-tie architecture is the result of optimization in the trade-off among robustness, fragility, resource limitation, and performance [1]. Polouliakh et al. and Yan et al. have shown that a narrow intermediate layer in the bow-tie architecture provides the classification capability of inputs [17] [22]. In terms of control theory, Wang et al. have proposed that bow-tie architecture possesses controllability [23], and in line with this idea, Kitano and Ni et al. have suggested that bow-tie architecture retains high evolvability, with a capacity for adjusting the outputs to the environment [1] [24]. Although these studies have shown the potential functions or advantages of bow-tie architecture, how this architecture evolved has not yet been elucidated.

Friedlander et al. have proposed an evolutionary mechanism of bow-tie architecture based on their simulation using a linear network model [2], in which the biological networks were modeled by a set of matrices describing the network structure (Fig 2A). They demonstrated that the bow-tie architecture emerges when (1) the mutations occur in a multiplicative manner, and (2) the network evolves toward a certain target in-out relation (i.e., the goal matrix; see Fig 2) that is expressed by a rank-deficient matrix. The condition (1) models the characteristics of biological mutations, which have been reported to be multiplicative [25] [26], and tend to minimize link intensity [27]. The condition (2) reflects biological inputs that are often redundant [28], such as image inputs in the retinal neural network [27] or Toll-like receptor signaling pathways [29]. Although Friedlander et al. [2] proposed a clear formulation for addressing the bow-tie evolution and presented a plausible hypothesis, it remains unclear how generalizable their hypothesis is, regardless of the simulation parameters.

**Fig 2.**
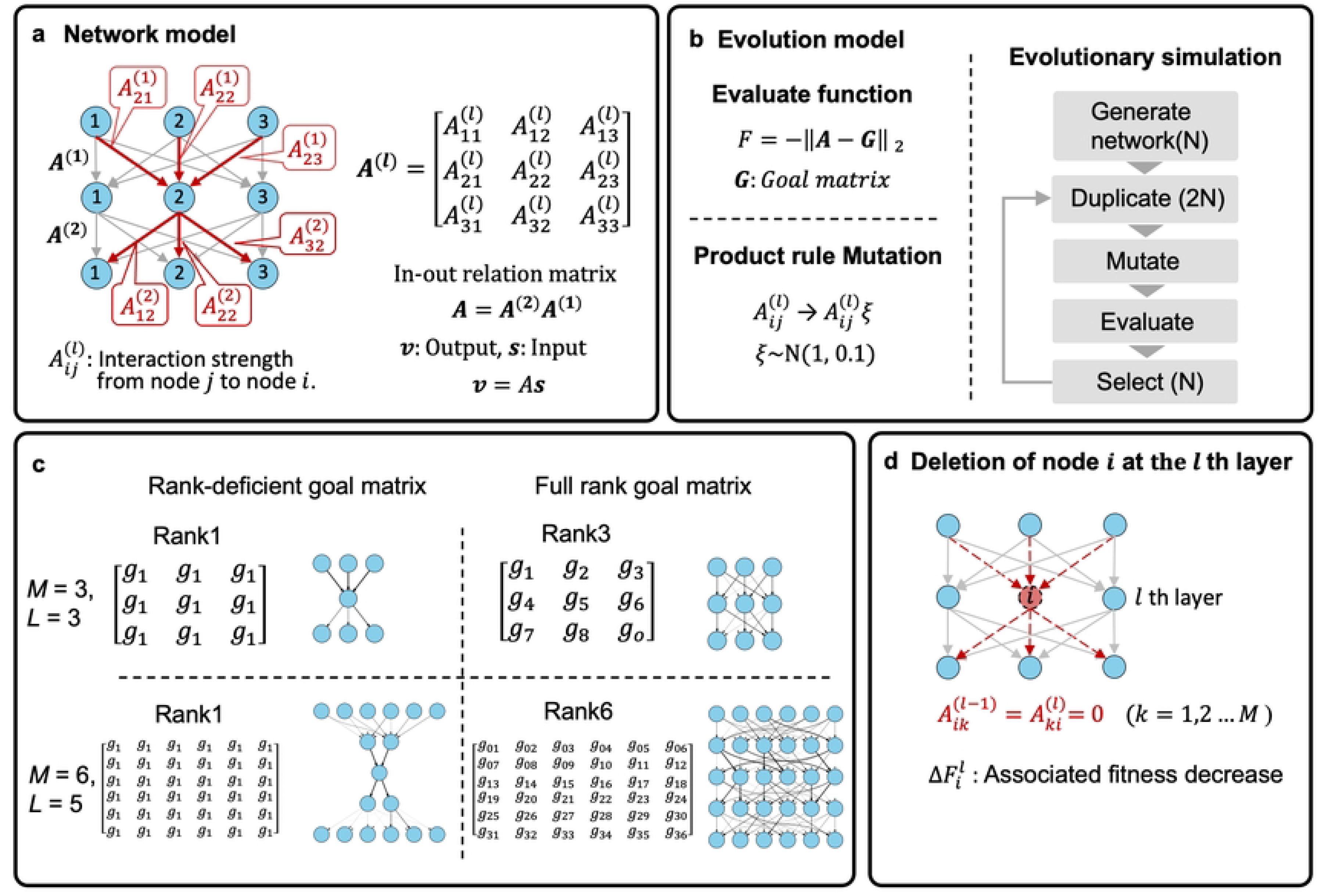
Evolutionary simulation of bow-tie architecture. (**a**) Network model. The network is modeled as a hierarchical linear network. All links between layers are described by a matrix. The *ij* element, 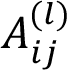, represents the strength of output from node *j* in layer *l* to node *i* in layer *l* +1. The product of these matrices produces the in-out matrix of network *A*. (**b)** Evolutionary model of the network. The evaluate function is defined by the distance between in-out matrix *A* and goal matrix *G*. ǁ*·*ǁ_2_ is the Frobenius norm. Mutation is modeled as the product of link intensity *A*^(*l*)^ and ξ generated from a Gaussian distribution. Evolution is simulated by repeating duplication, mutation, evaluation, and selection. (**c)** A different rank of goal matrix is randomly generated. The typical evolved network under the given goal matrix is shown next to the goal matrix. Bow-tie architecture emerges under the rank deficient goal matrix. (**d)** Deletion of node *i* at the *l* th layer. The effect of node *i* is evaluated by assessing the fitness decrease upon removal of node *i*. Removal of node *i* is executed by 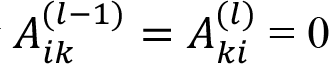.

In this study, we report that their hypothesis is only valid when the link intensities representing molecular interactions are small at the initial condition of the evolutionary simulation. This indicates that the network structure at the beginning of the evolution plays a crucial role in the emergence of bow-tie architecture. Interestingly, when evolution starts from weak initial link intensities, the bow-tie architecture transiently appears even in the absence of the previously proposed condition (2). In the present study, we analyze the mechanism of the emergence of bow-tie architecture by using a simple ODE model. Based on the identified mechanism of emergence of bow-tie architecture, we further identify natural conditions under which the bow-tie architecture evolves independently of such artificial parameters as the initial link intensities and goal matrix rank.

## Model

To investigate the evolution of the bow-tie architecture, we consider layered feedforward networks with *M* nodes in each layer. We adopted the linear network model proposed by Friedlander et al [2], in which link intensities from the *l* th layer to the *l* + 1 th layer are described by the *M* × *M* matrix ***A***^(***l***)^ (Fig 2A). The *ij* element, 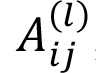, is the link intensity from node *j* in the *l* th layer to node *i* in the *l* + 1 th layer and is defined as 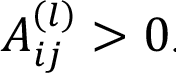. When an input signal ***s*** is applied to the network with *L* + 1 layers, output vector ***v*** is obtained as the product of matrices *A****s*** = ***v***, where *A* is the in-out relation matrix of the network defined by *A* = *A*^*L*^*A*^*L*–1^. . . *A*^1^ (Fig 2A). The network evolves toward an ideal in-out relation matrix ***G*** (hereinafter referred to as the goal matrix) (Fig 2B). We adopted a randomly generated goal matrix with different ranks (Fig 2C). The evaluation function, *i.e.*, fitness, is defined by *F* = −‖*A* − *G*‖ _2_, where Frobenius norm ‖·‖ _2_ represents the sum of the square of all matrix elements, and thus -*F* denotes the distance between the in-out relation matrix ***A*** and the goal matrix ***G***. We evolved the network by optimizing the in-out relation matrix ***A*** so that fitness *F* is maximized through the following genetic algorithm (Fig 2B). We generate *N* individuals having an *L* + 1 layered network represented by a set of matrices *A*^1^, …, *A*^*L*^. In each generation during evolution, the individuals are duplicated so that the population size is 2*N* and mutations are randomly introduced into 20% of individuals in that population. For mutations, we follow Friedlander et al., assuming the product rule mutation [2]; a randomly selected network link 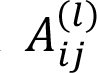 is altered as 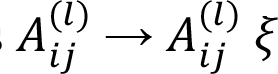 with a random number *ξ* generated from *N(1,0.1), i.e.,* a Gaussian distribution with mean one and variance 0.1. Then, the fitness *F* in 2*N* individuals is evaluated, and finally the top *N* individuals are selected by tournament selection with group size four [2] [27] [30]. Evolution is simulated by repeating these processes (Fig 2B). The individuals are considered to be fully evolved when their average fitness value reaches -0.01 or higher, and the network of individuals with the highest fitness *F* is analyzed. Throughout this study, the population size *N=100* is used. Whether network architecture is bow-tie or not is judged by calculating the decrease in fitness associated with the removal of each node. Deletion of node *i* at the *l* th layer is implemented by setting 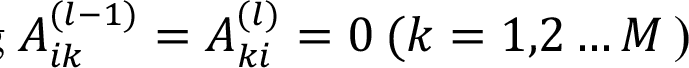, and the absolute value of the associated fitness decrease is denoted by 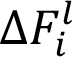 (Fig. 2D). The ratio of fitness decrease 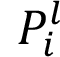 is then estimated as 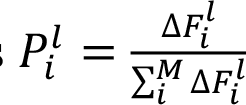. We define an active node as a node whose removal causes a decrease in fitness satisfying a criterion 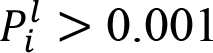. When the number of the active nodes in a middle layer (i.e., *l* = 2 – *L*-1) is smaller than *M*, the network is defined as a bow-tie network.

## Results

### Bow-tie networks evolve only when the initial link intensities are small

First, we attempted to reproduce the simulation of Friedlander et al. [2] by using the 6 nodes x 5 layers network, i.e., *M* = 6 and *L* = 4 (Fig 3A). We started the evolutionary simulation from a random configuration of 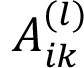 with a small initial link intensities *A*_0_ = 0.01, which is defined by the Frobenius norm of the initial in-out relation matrix ‖*A*‖ _2_. We were able to reproduce the previous results [2], confirming that the network evolves to the bow-tie architecture when the goal matrix is rank deficient (Fig 3A). Although we adopted tournament selection following the previous result, in a separate analysis we obtained almost the same result by adopting elite selection (Fig S1). Figure 3A illustrates the most frequent number of active nodes for each layer. We here refer to a middle layer with the smallest number of active nodes as a "waist" and denote its number of active nodes as the "waist width". The waist width decreased as the rank of the goal matrix decreased; for a low rank goal matrix, the network with a narrow waist width evolved, while a network with a waist width = M appeared for the full rank goal matrix (Fig 3A).

**Figure 3.**
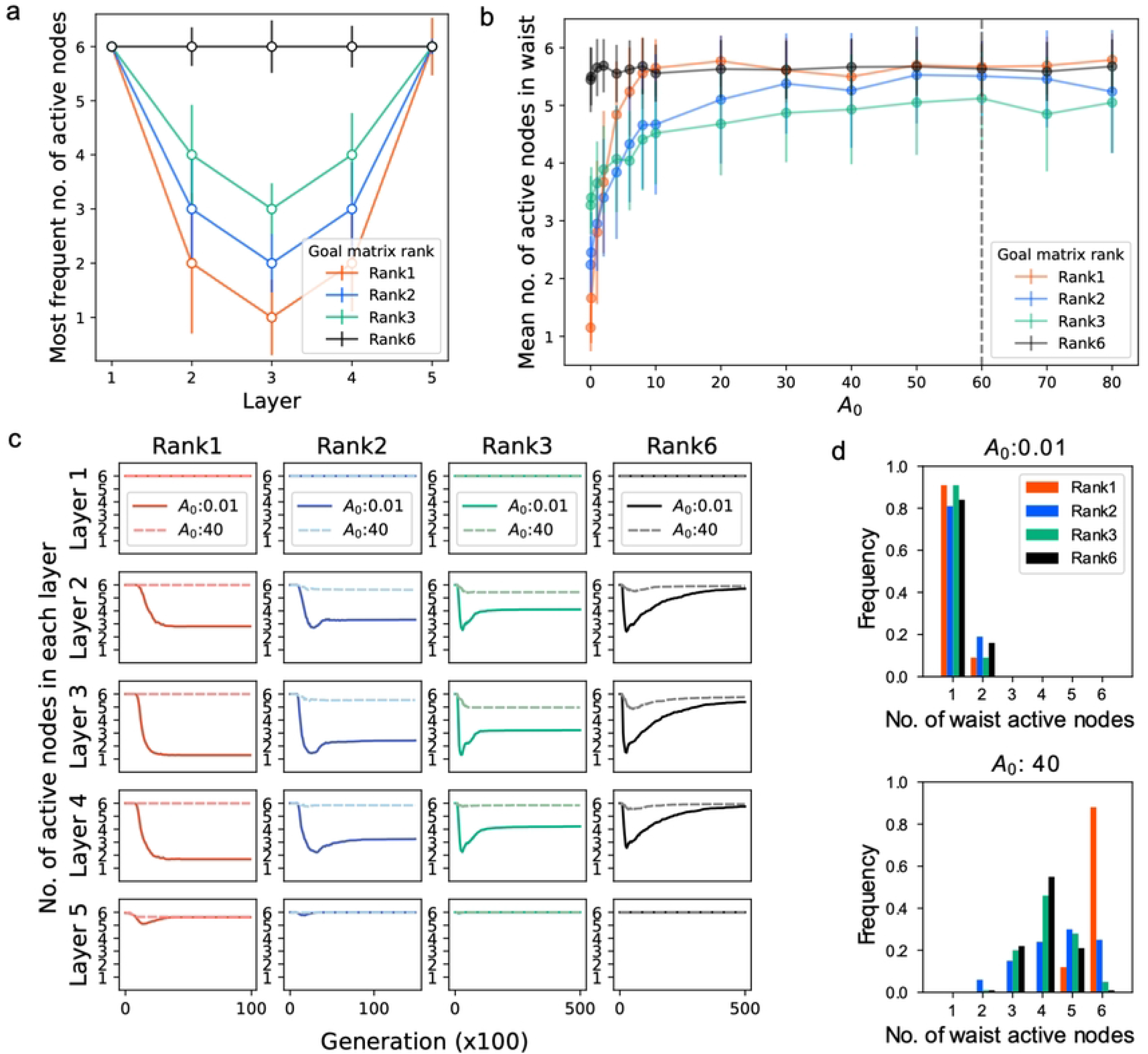
Simulation results with a 6 nodes x 5 layers network. (**a**) Reproduction of the previous results [2]. The Y-axis shows the mode among 100 runs of the number of active nodes in the most-adapted network. Initial link intensity is set to *A*_0_: 0.01. The error bars represent the std. Goal matrix elements are randomized under the conditions of rank1 (red), rank2 (blue), rank3 (green), and rank6 (black). The norm ‖***G***‖_2_ is normalized to the same value (‖***G***‖_2_ = 60). (**b)** The bow-tie emergence crucially depends on the initial link intensity. X-axis: the link intensity of the initial network. Y-axis: Mean number of active nodes in the waist layer in an adapted network among 100 runs. The error bars are the std. The dashed line shows ‖***G***‖_2_ = 60. (**c)** Evolution trajectories of the number of nodes in each layer. Simulation starts from *A*_0_ = 0.01 in the solid lines and *A*_0_ = 40 in the dashed lines. Trajectories are averaged among independent simulation runs (*n=100* for each color). The simulation runs that reach *F >* -0.01 are used. (**d)** Distribution of the narrowest waist width that the network experienced during evolution (*n=100* for each color).

In contrast to the previously reported result [2], when starting from a bit larger initial link intensities *A*_0_, this emergence of the bow-tie architecture vanished even for the low rank goal matrix. We simulated the evolution of a network starting from various values of *A*_0_. Figure 3B shows the waist width of the evolved network against *A*_0_, in which each dot is the average value of over 100 runs, and illustrates that the bow-tie architecture evolves only at a range of *A*_0_(0.1∼10). This range is quite narrow compared with the Frobenius norm of the goal matrix, i.e., 60, denoted by the black dashed line in Figure 3B. When we started the simulation with an *A*_0_ larger than 10, the network did not converge to a bow-tie structure in most cases (specifically for rank = 1).

The evolutionary dynamics also exhibited a remarkable dependence on initial link intensity *A*_0_. Figure 3C shows the time series of the number of active nodes in each layer for a large (= 40) and small (= 0.01) *A*_0_ with different goal ranks. When the goal matrix was full-ranked, the number of the active nodes in the 2–4th layers transiently decreased and then converged to 6 for the small *A*_0_ (Fig 3C; Rank 6, solid black line), indicating a transient emergence of the bow-tie architecture. For the large *A*_0_, the number of active nodes did not show such a transient drop (Fig 3C; Rank 6, dashed black line). When the goal rank was low, the network starting from the small *A*_0_ converged to the bow-tie network (Fig 3C; Rank 1, Rank 2, and Rank 3, solid line), while the network starting from the large *A*_0_ did not show even a transient emergence of the bow-tie architecture (Fig 3C; Rank 1, Rank 2, and Rank 3, dashed line). Figure 3D shows the distribution of the narrowest waist widths that the network reached during the course of evolution, and indicates that the network evolves to bow-tie architecture regardless of the rank of the goal matrix if it starts from a small *A*_0_ (Fig 3D; upper panel), while bow-tie architecture could not be produced even in the low rank goal matrix when the network started from a large *A*_0_ (Fig 3D; lower panel). These results suggest that the appearance of the bow-tie architecture in the early phase of evolution is determined by the initial link intensities *A*_0_, rather than the rank of the goal.

### Analysis of bow-tie architecture evolution by the dynamical system

To determine the reason for the dependence on the initial link intensities *A*_0_, we consider a phenomenological ordinary differential equation (ODE) of the network 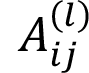 that qualitatively mimics the evolutionary simulation described above. Based on the gradient descent, the time evolution of link intensity 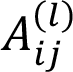 is given by

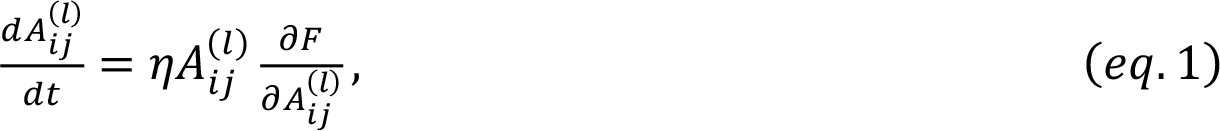

where the product-rule mutation is implemented by the learning rate proportional to 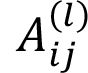. This optimization algorithm, which we refer to here as the multiplicative gradient descent, describes the multiplicative changes in the link intensity, and therefore the larger link changes more drastically. The learning rate *η* is set to a constant value *η* = 10^-4^. We confirmed that the results obtained using the ODE model were almost the same as those by the original model (Fig S2)—namely, the ODE model reproduced bow-tie architectures for a rank-deficient goal matrix *G* and a small *A*_0_. We also confirmed that the ODE model exhibited a dependence on *A*_0_similar to that of the original model (Fig S2A and B). The ODE model also recapitulated the transient appearance of the bow-tie in the original model (Fig S2 and D).

Using this ODE model, we here analyzed how the bow-tie architecture emerges in the early stage of the time evolution when starting from a small *A*_0_. For simplicity, we consider the networks that consist of 3 layers with 2 nodes each (Fig S3A) with the simplest rank-1 goal matrix G defined as

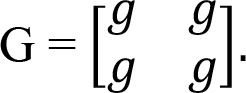

The following argument can also be generalized for an arbitrary G with rank-1 or rank-2 (the Supplemental materials). Under this condition, the evaluation function is given as follows (eq. 2):

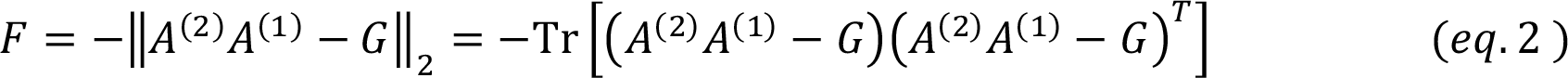

The adaptation process is modeled as the multiplicative gradient descent eq.1 as follows (see the

Supplemental materials for details):

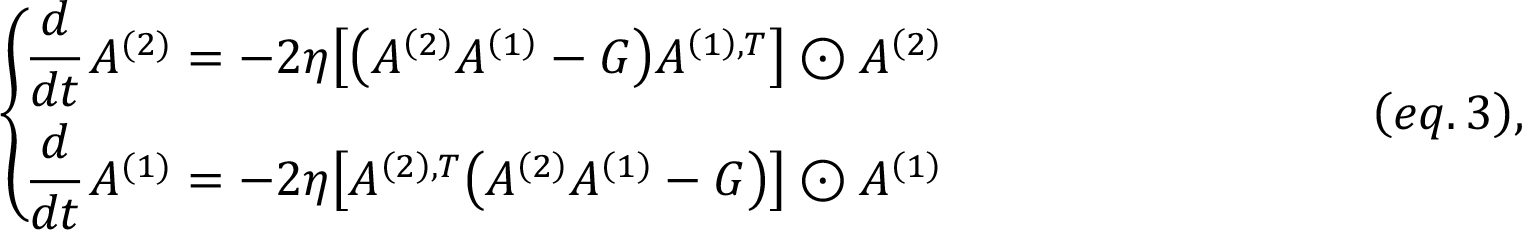

where ⊙ is the Hadamard product, *i.e.*, (*A* ⊙ *B*)_*i*j_ = *a*_*i*j_*b*_*i*j_, and *A*^(*i*),*T*^is the transposed matrix of *A*^(*i*)^. Since we consider the early stage of the dynamics starting from *A*^(1)^and *A*^(2)^ that are much smaller than *G*, the term N*A*^(2)^*A*^(1)^ − *G*O can be approximated as −*G*, and thus eq. 3 is simplified as follows:

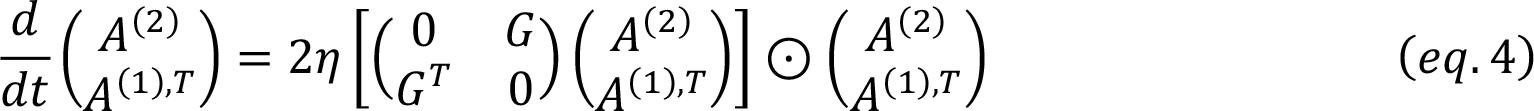

By expanding eq. 4, the following equation is obtained:

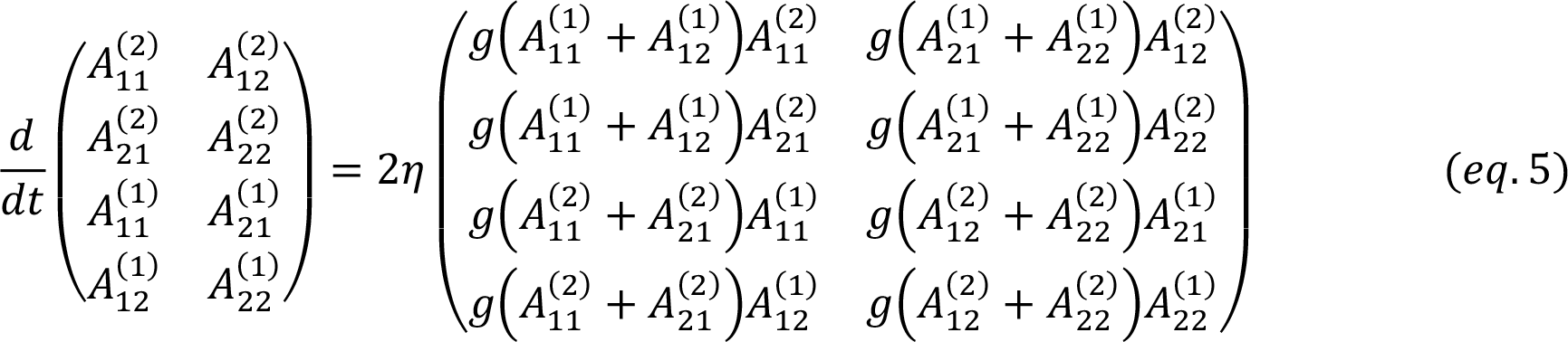

Note that the first and second columns are independent because the time evolution of 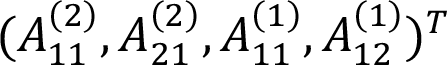 does not depend on the 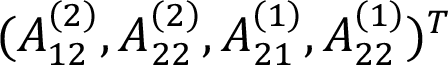 The first column of eq. 5 is given as

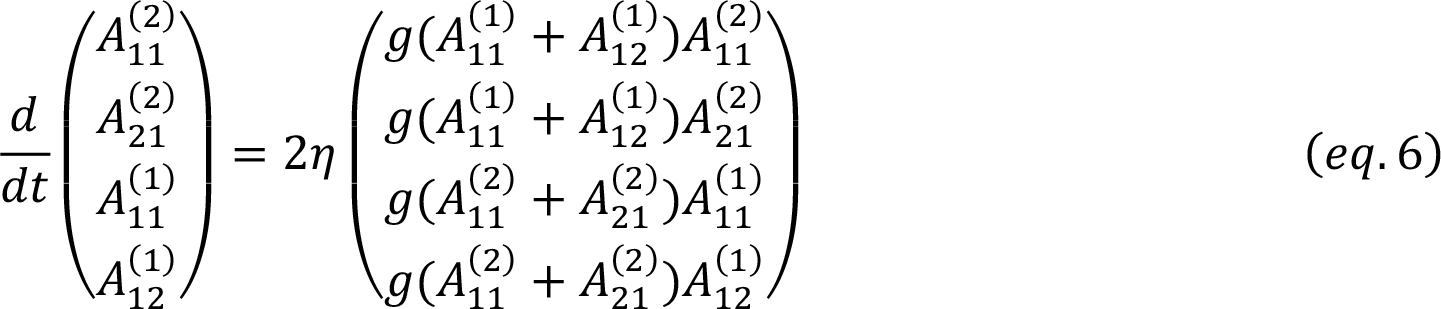

Here, we define *l*_*k*_ and *R*_*k*_ as follows.

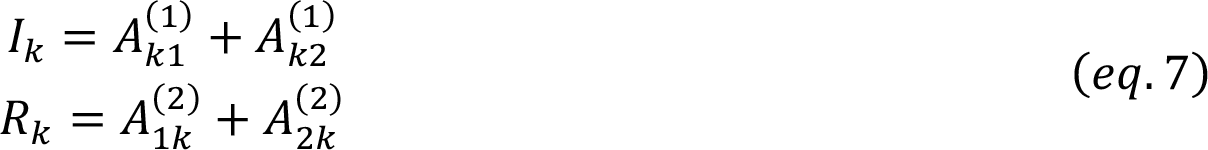

*l*_*k*_ represents the sum of input strength to the node *k* in the middle layer, whereas *R*_*k*_ denotes the sum of output strength from node *k* in the middle layer (Fig S3A). From eq. 6 and eq. 7, the following relation holds:

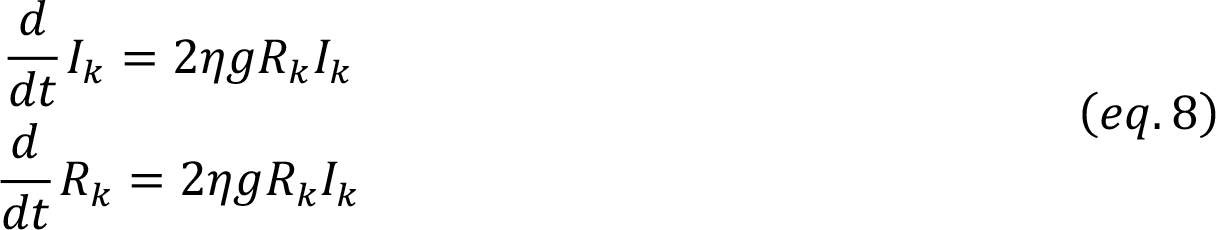

Thus, 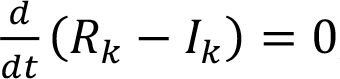, indicating *R*_*k*_ − *l*_*k*_ = *R*_*k*,0_ − *l*_*k*,0_, where *R*_*k*.0_ and *l*_*k*,0_ are the initial values of *R*_*k*_ and *l*_*k*_ (Fig. S3B). Substituting this expression into eq. 8, 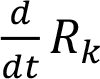 is described as follows.

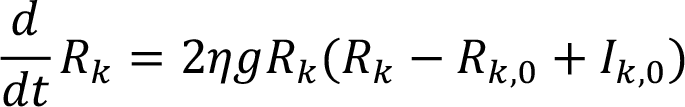

By solving this, the time evolution of *R*_*k*_ is derived.

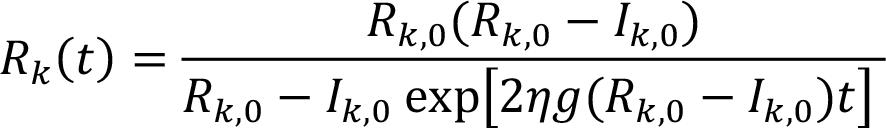

This indicates that *R*_*k*_ diverges within a finite time, 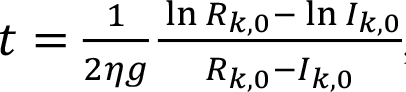, and *l*_*k*_ also diverges since *R*_*k*_ − *l*_*k*_ = *const*. The same argument holds for the second column of eq.5, and thus one of the two columns diverges prior to the other, which demonstrates the emergence of bow-tie architecture (see Fig. S3B). For the case of *R*_*k*,0_ ≈ *l*_*k*,0_, the divergent time is written as *t*∼1/*gR*_*k*,0_ (see the Supplemental materials), which indicates that the column that has a larger *R*_*k*,0_ diverges first.

Although the rank-1 goal matrix with 2 nodes is assumed here, the independent relationships between columns of 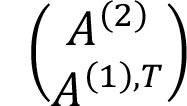 in eq. 4 hold, regardless of the goal matrix rank and number of nodes (see the Supplemental materials). Therefore, the bow-tie emergence with the divergence of *R* and *I* for one of the columns always occurs if the approximation *G* − *A* ≈ *G* is valid. The intuitive explanation for this effect is as follows: In the case of *G* − *A* ≈ *G*, all 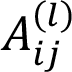 grow to increase the fitness, but the multiplicative nature of the growth leads to the significant difference 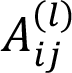 among the *l* th layer, *i.e.*, 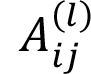 starting from the smaller value grows more slowly, which causes the bow-tie architecture. This is not the case for the condition under which the system starts from a large initial value *A* ≈ *G*, because the fitness reaches the optimal value before the product rule mutation causes a significant difference of 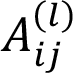. These arguments help to explain how the bow-tie architecture emerges in the early phase of evolution when the initial link intensities are small, regardless of the goal rank.

### The environmental fluctuation facilitates the emergence of the bow-tie architecture

The aforementioned results assume a fixed goal matrix, which corresponds to the evolution under a fixed environment. However, in a more realistic situation, the element of the goal matrix (ideal in-out relation) fluctuates due to the environmental fluctuation or mutations in the numbers of the receptors and downstream genes. To examine the effect of an environmental fluctuation on bow-tie architecture evolution, we introduced the variable goal matrix without changing the dimension, rank, or norm. First, the network evolves under the first goal matrix for 150,000 generations, which is enough time for the fitness to reach 0.01. Second, the goal matrix is altered to the next one. We found that the network architectures are maintained irrespective of any such change in the goal matrix during evolution; the waist width did not change before and after the change in goal matrix (Fig S4), indicating that the waist width is not affected by the change in goal matrix but determined by the *A*_0_. The number of active nodes transiently increases immediately after the change of goal matrix. This is attributed to the definition of active nodes; due to the decrease in fitness associated with the change in goal matrix, the difference of 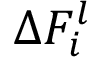 (i.e., the fitness decrease by the *i* th node removal) among nodes in the *l* th layer is less clear, and thus the value of 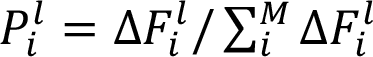 tends to exceed the threshold, which leads to an increase in the waist width. These results reflect that the bow-tie architectures are robust to environmental fluctuations that do not alter the rank and norm once the bow-tie architecture emerges. Consistent with such a conclusion, after the network adapts to the initial goal, the network architecture does not change by repeated changes in the goal matrix. Figure 4A shows the time-course of the active nodes in each layer during the repeated goal change with a fixed goal rank every 1000 generations, indicating that the network architecture is maintained. Interestingly, after the adaptation to the goal matrix, the network structure is still maintained even for the changes in rank of the goal matrix. The network is first evolved to adapt to the rank-1 goal, and then the goal matrix is repeatedly switched to the randomly generated full-rank goal matrix every 10^3^ generation (Fig 4B). This procedure did not reshape the bow-tie architecture during 10^4^generations.

**Figure 4.**
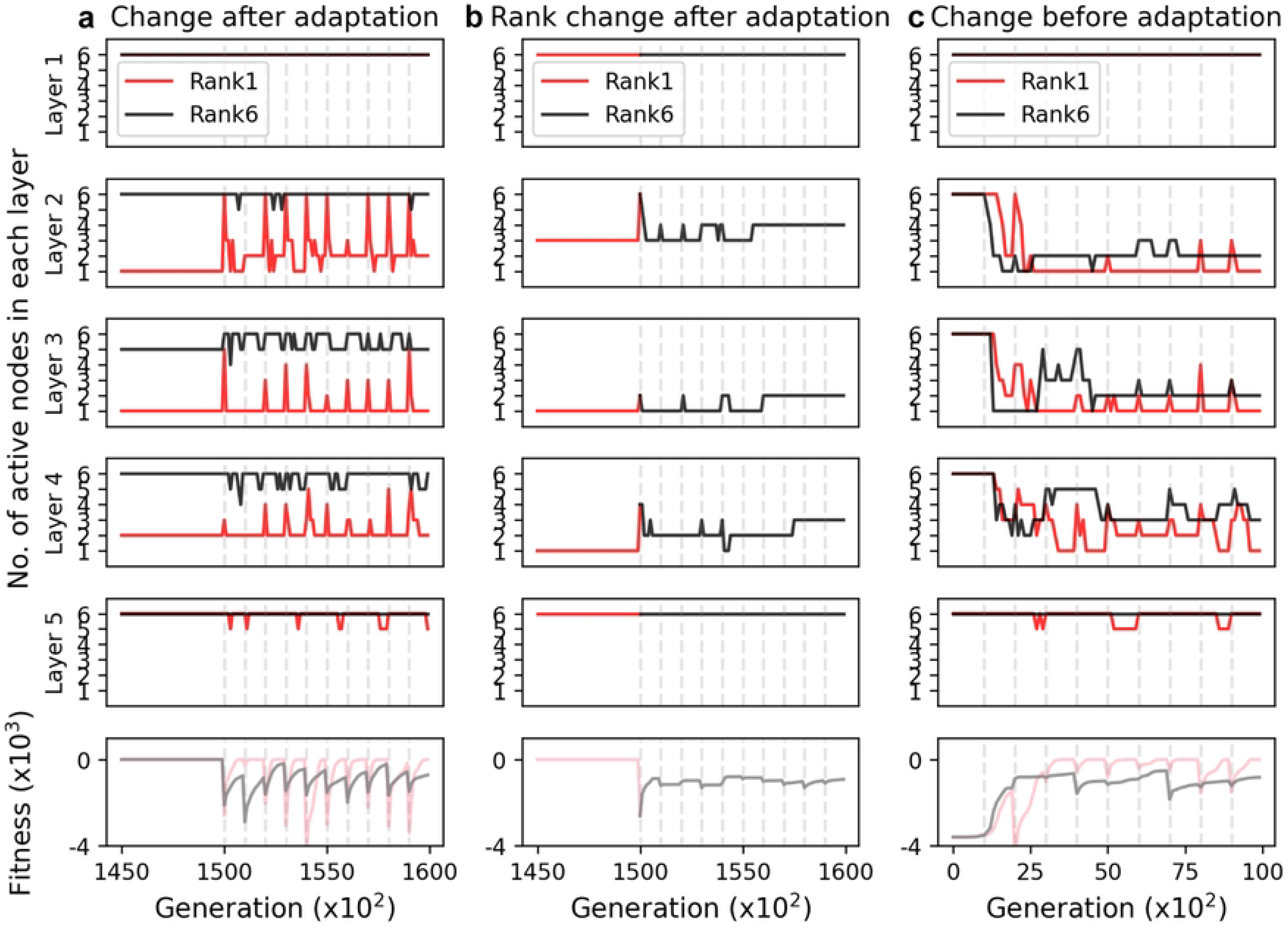
Introduction of goal matrix fluctuation in a network with M=6 L=4 (5 layer). The goal matrix is changed every 1000 generations (dashed line) after or before the adaptation. The number of nodes in each layer (upper 5 panels) and the fitness of the most-adapted network in the population (the bottom panel) are plotted against the generation. (**a)** Sequential goal matrix change after fitness is converged. The goal matrix is changed without changing the norm and rank (red: rank1; black: rank6). (**b)** Sequential goal matrix change after fitness is converged. The goal matrix rank is changed from rank1 to rank6 in the first change. (**c)** Sequential goal matrix changes from the beginning of evolution. The goal matrix rank and norm do not change (red: rank1; black: rank6).

Although these data indicate that the environmental fluctuation does not affect the previously emerged bow-tie architecture, this is not the case when the fluctuation takes place at the beginning of the evolution. When starting from a small initial link intensities *A*_0_ (*A*_0_ = 0.01), the initial random network develops the bow-tie architecture by repeated fluctuation of the goal matrix, regardless of the goal rank (Fig 4C). This can be understood by the fact that the time-averaged goal matrix can be effectively interpreted as a rank-1 matrix, which facilitates the bow-tie emergence.

### The gene duplication accelerates the emergence of the bow-tie architecture

Finally, we examined the effect of expansion of the goal matrix. During evolution, the number of the input and output nodes can be increased by the gene duplication, and by de-novo addition (emergence) of the receptor molecules and downstream genes. Therefore, we investigated whether the non-bow-tie network switches to the bow-tie architecture in response to an increase in the goal matrix dimension. We found that an increase in the goal matrix dimension indeed leads to the emergence of the bow-tie architecture even when starting from a large link intensity (Fig 5). The details of the simulation procedure are shown in Fig 5A: Evolutionary simulations start from the 3 nodes by the 3 layers network during the 1000th generation. Then, the goal matrix is expanded from a 3×3 matrix to a 9×9 matrix at the 1000th generation (Fig 5A). Consequently, the network converged (|Fitness| < 0.01) to the bow-tie architecture after expanding the goal matrix (Fig 5B and Fig 5C). Note that although the input and output nodes are increased by the goal matrix expansion, the evolved waist width is lower than that before expanding the goal matrix (Fig 5C and 5D). This would also be explained from the proposed scenario starting from a small link intensity; the expansion of the goal matrix results in the increase in the norm of the goal matrix *G*, causing ‖*G*‖_2_ ≫ ‖*A*‖_2_, and subsequently leads to evolution of the bow-tie architecture as discussed in the above section.

**Figure 5.**
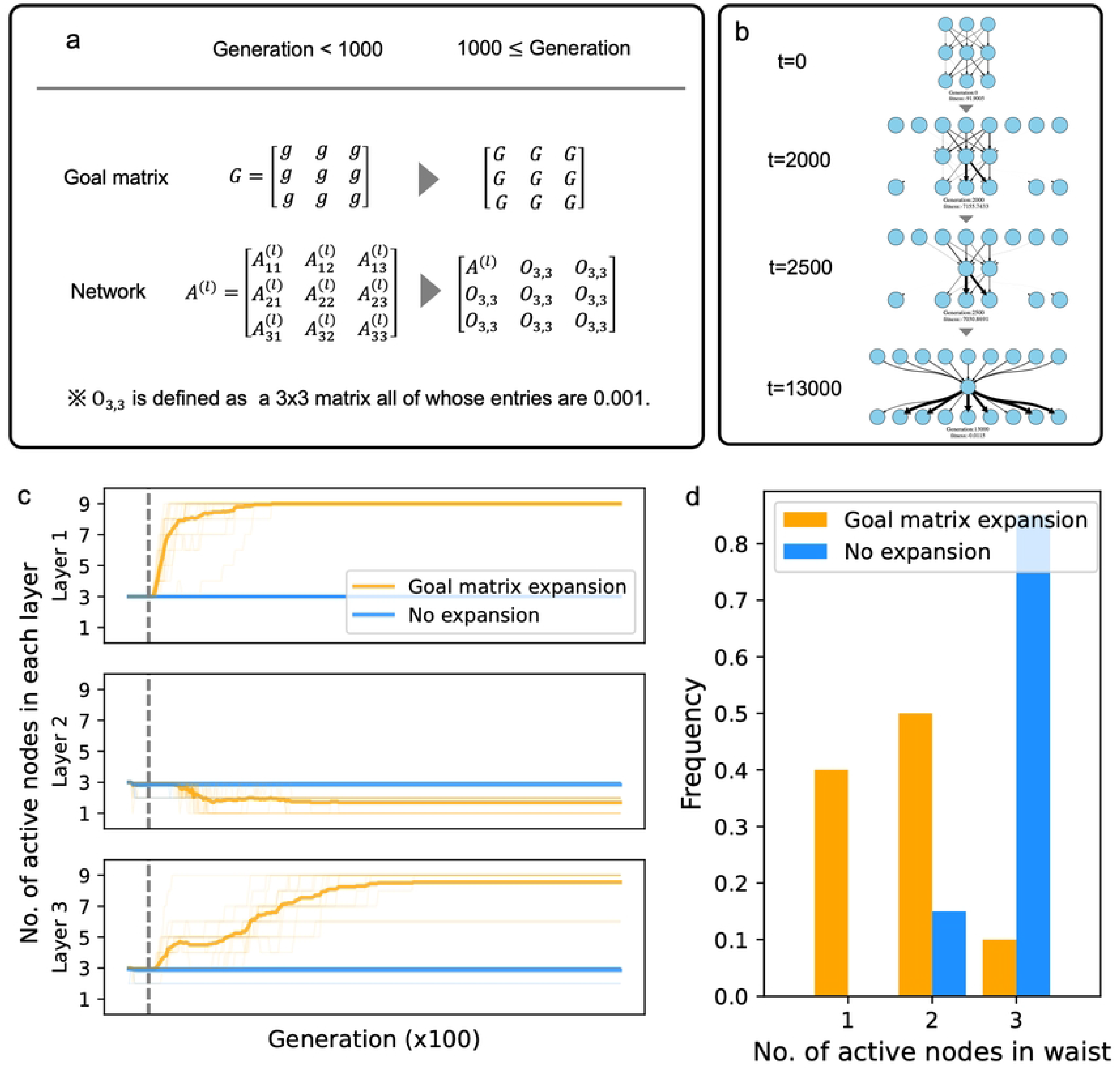
Expansion of the goal matrix drives evolution of the bow-tie architecture. a) The procedure of goal matrix enlargement. (**a**) The 3×3 goal matrix G is expanded to the 9×9 matrix at the 1000^th^ generation. In association with the goal matrix expansion, the network matrix ***A***^(***l***)^ is also expanded to a 9×9 matrix by padding using the near zero 3×3 matrix O_3,3_. **(b)** An example of the network during evolution after the goal expansion. (**c)** Evolutionary trajectories of the network. X-axis: Generation. Y-axis: Number of nodes in each layer. The initial norm *A*_0_is 10. The orange line represents the case where the 3×3 goal matrix with rank 1 is expanded to 9×9 at the 1000 generation (dashed line), while the blue line represents the simulation with no enlargement. (**d)** Statistics of the number of active nodes in the adapted network for each layer (x-axis) with (orange) or without (blue) the goal matrix expansion.

## Discussion

In this study, we investigated the conditions under which the bow-tie architecture emerges by using an evolutionary simulation with a linear network model. Although the previous study [2] suggested that bow-tie architecture evolves by (1) the product rule mutation with (2) a rank deficient goal (Fig 2), our extensive simulations under various initial conditions revealed that these conditions (1) and (2) are not sufficient for the appearance of bow-tie architecture, and that an additional condition is required—namely, the norm of the initial in-out relation matrix *A*_0_ must be much smaller than *G*. The evolution from small *A*_0_(*i.e.,* small link intensity) may correspond to the early stage in the evolution of a signaling network, where molecular interactions are almost entirely absent. Consistent with such an interpretation, a core of the bow-tie architecture is well conserved (e.g., the G protein), implying that it has a very old origin [19] [31] [32].

As discussed for the ODE model, the emergence of bow-tie architecture in the situation *A*_0_ ≪ *G* (i.e., *G* − *A* ≈ *G* in eq. 3) is attributed to the multiplicative nature of the growth in 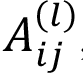, which can cause the significant difference in 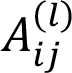 among the *l* th layers. In addition, we demonstrated that bow-tie architecture transiently emerged regardless of the goal matrix rank when the initial link intensities were small (Fig 3C and 3D). This indicates that the small initial link with the multiplicative mutation is essential for the emergence of bow-tie architecture, even it is transient, and the rank deficient goal matrix only determines whether or not the transient bow-tie architecture is maintained for a long time.

In a more realistic situation, the ideal in-out relation (goal matrix) can fluctuate. We confirmed that, once the bow-tie architecture is established, it is robust against the goal matrix fluctuations (Fig 4B and 4C). This result is consistent with the previous suggestion that bow-tie structures are maintained under perturbation and fluctuation [1] [4] and with the well-conserved core of actual bow-tie architecture [19] [31] [32]. The fluctuation even enhances the emergence of bow-tie architecture at the early stage of the evolution (Fig 4C). This phenomenon does not depend on the goal matrix rank. These results may explain the ubiquitous appearance of the bow-tie network [1].

Figures 3B and C show that a network starting from strong links (*i.e.*, a large A_0_) cannot evolve into a bow-tie architecture. However, we demonstrated that the bow-tie architecture appears even from a large link in the presence of large-scale perturbations such as genome duplications. The simulation in Figure 5 mimics the increases in inputs (receptors) and outputs (downstream genes) resulting from an expansion of the goal matrix. From the analysis in Figure 5, we found that a rapid increase in the number of inputs and outputs narrowed the waist width in the network, leading to the bow-tie architecture. This suggests that the addition of inputs and outputs promotes the emergence of bow-tie architecture. A similar result was reported by Bauer et al., who found that a hub emerged when the number of nodes in the network was increased exponentially [33]. In fact, in the GPCR signaling system, which forms a narrow waist bow-tie architecture, the number of receptors and downstream genes has been reported to increase rapidly during the evolution [31] [34]. However, further analysis will be needed to determine whether such expansion occurs after or before the formation of the bow-tie architecture.

Our hypothesis would suggest that the bow-tie architecture emerges as a byproduct of evolution, rather than as a result of the evolutionary adaptation. However, it does not deny the functional advantages of the bow-tie architecture suggested by the previous studies [17] [22] [23] [24]. For example, the bow-tie architecture can be controlled by controlling input signals [23], and it shows fast adaptation to novel environments and high evolvability [24]. Such fast adaptation of the bow-tie architecture was also seen in our simulation (Fig S4A, red lines). We also found that network generally evolves to bow-tie architecture under the fluctuating in-out relation (fig. 4). This suggests that the bow-tie structure can adapt diverse inputs and outputs, and thus providing generalization capability as claimed in previous studies [17] [22]. On the other hand, the bow-tie network has a high lethality to perturbation to the core nodes [35] [36] and is disadvantageous for use in a search for alternative core genes [4]. The bow-tie architecture would have been selected as the result of a trade-off between these advantages and disadvantages [4]. Nonetheless, the advantages of the bow-tie architecture should contribute to the long-term conservation of bow-tie structures.

A more intuitive mechanism of evolution of bow-tie architecture includes the cost for maintaining link intensity 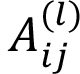 When the cost of the L1 regularization term 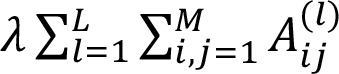 is included in the fitness, the networks evolve towards the bow-tie architecture (Fig S5) independent of the initial link intensities A_0_, while the cost of the L2 regularization term 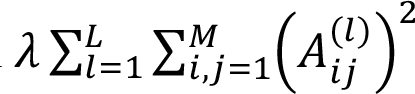 enhance the bow-tie structure (Fig S5). These results are explainable from the well-known effect in the data sciences [37]that the L1 regularization term reduces the network links. However, no biologically plausible explanation can be made for the introduction of the L1 term rather than the L2 term, and thus we concluded this assumption would be artificial.

In summary, we suggested two evolutionary scenarios to explain the emergence of the bow-tie architecture: 1) the bow-tie architecture emerges in the early stage of evolution and 2) the bow-tie architecture emerges immediately after the addition of inputs and outputs. Both scenarios are based on a mathematical condition: A_0_ << G (large gap between initial and goal link intensities). Although the linear feedforward network was adopted here, a more realistic interaction including nonlinearities and feedback loops should be examined in the future. Further, a bioinformatic analysis revealing the origin and evolutionary history of bow-tie architecture will be required to support our hypothesis.

## Acknowledgements

We thank all members of the Aoki Laboratory for the helpful discussion. This study was supported in part by Cooperative Study Program of Exploratory Research Center on Life and Living Systems (ExCELLS; program No. 21-102, 22-102 to N.S.).

## Author contributions

T.I., Y.K., K.A. and N.S. designed research. T.I. performed research and analyzed data. T.I. and N.S. constructed mathematical model. T.I. and N.S. drafted the manuscript. All authors approved the final manuscript.

## Figures

**Figure S1.**
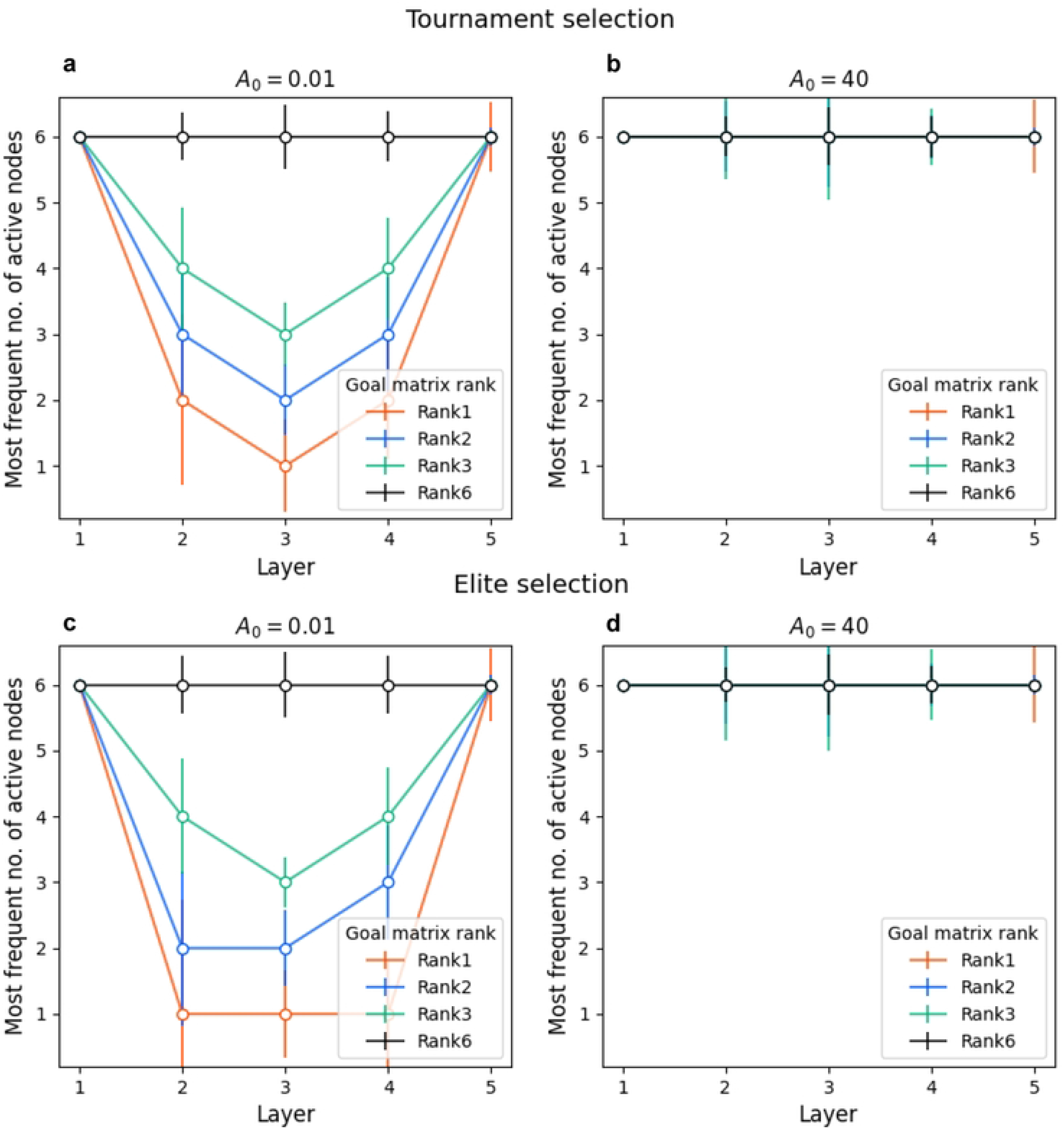
Comparison of evolutionary simulation by tournament and elite selection. The Y-axis shows the mode among 100 runs of the number of active nodes in the most-adapted network. (**a), (b)** Evolutionary simulation with tournament selection starting from (**a)** small initial intensity and (**b)** large initial intensity. (**c), (d)** Evolutionary simulation with elite selection starting from (**c)** small initial intensity and (**d)** large initial intensity. The error bars are the std. The top left panel (a) is identical to Fig. 3a.

**Figure S2.**
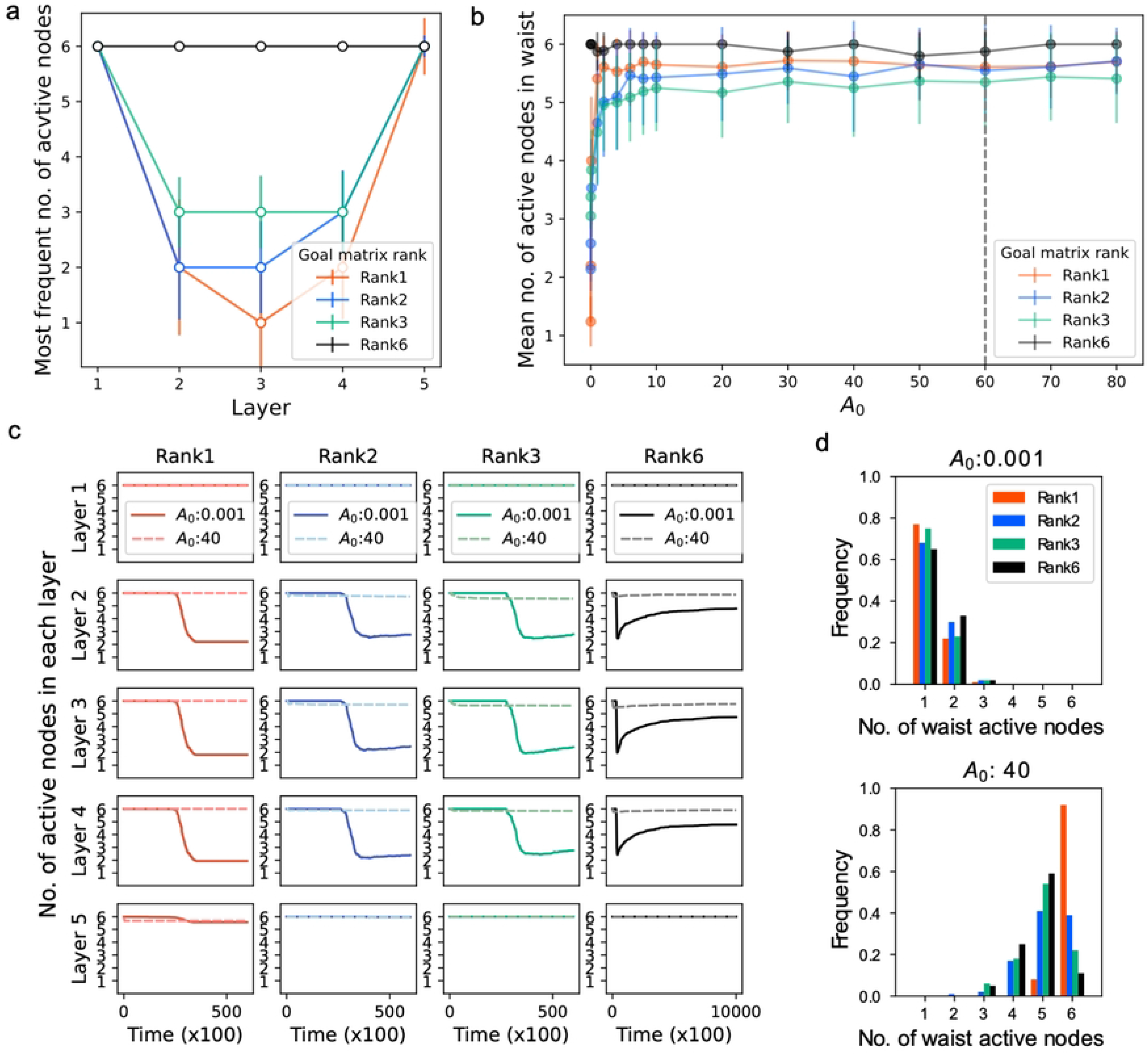
Simulation results with a 6 nodes x 5 layers network in the ODE model. The numerical integration is performed by the fourth-order Runge-Kutta method. (**a)** Reproduction of the previous results [2]. The Y-axis shows the mode among 100 runs of the number of active nodes in the most-adapted network. The number of runs is [rank1: 100; rank2: 100; rank3: 99; rank6: 5]. Initial link intensity is set to *A*_0_: 0.01. The error bars represent the std. Goal matrix elements are randomized under the conditions of rank1 (red), rank2 (blue), rank3 (green), and rank6 (black). The norm ‖***G***‖_2_ is normalized to the same value (‖***G***‖_2_ = 60). (**b)** The bow-tie emergence crucially depends on initial link intensity. X-axis: the link intensity of the initial network. Y-axis: Mean number of active nodes in the waist layer in an adapted network among 100 runs. The error bars are the std. The dashed line shows ‖***G***‖_2_ = 60. (**c)** Evolution trajectories of the number of nodes in each layer. Simulation starts from *A*_0_ = 0.001 in the solid lines and *A*_0_ = 40 in the dashed lines. Trajectories are averaged among independent simulation runs (*n=100* for each color). The simulation runs that reach *F >* -0.01 are used. (**d)** Distribution of the narrowest waist width that the network experienced during evolution (*n=100* for each color).

**Figure S3.**
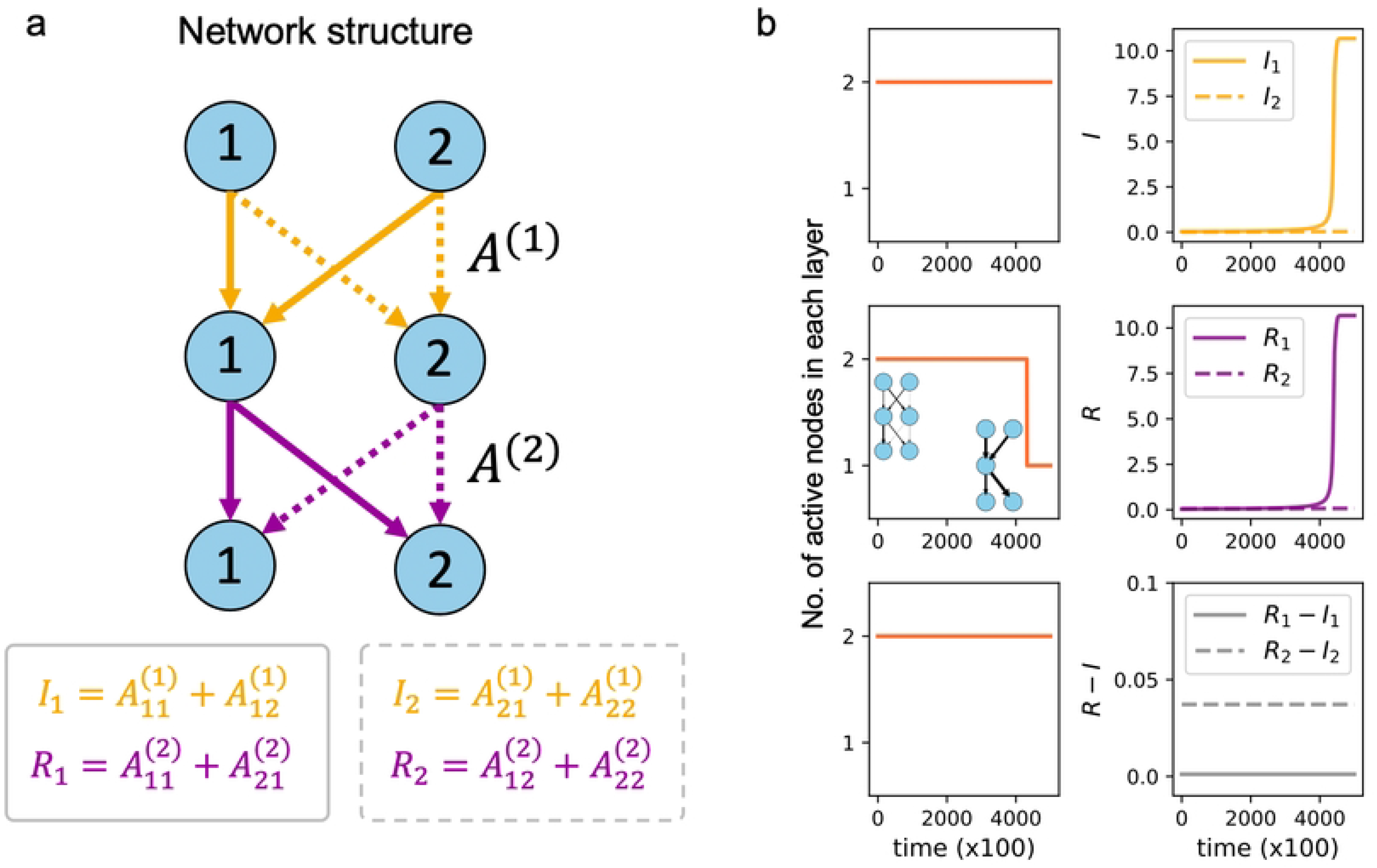
Analysis based on the ODE model with 2 nodes x 3 layers. **(a)** A 2×3 layer network described by 2 matrices A^(1)^ and A^(2)^. The links denoted by the thick lines (*l*_1_: yellow; *R*_1_: purple) are evolved independently of the dashed lines (*l*_2_: yellow; *R*_2_: purple). Each pair of *l*_*i*_ and *R*_*i*_ is interdependent by the constraint of *R*_*i*_ – *l*_*i*_= const when ***A*** ≪ ***G***. (**b)** Evolution toward the rank1 goal matrix (‖***G***‖_2_ = 60) from the initial norm *A*_0_ = 0.01. Left panel: Evolutionary trajectories of the network. Y-axis: Number of nodes in each layer. The initial norm *A*_0_ is 0.01. Right panel: Evolutionary trajectory of *I, R,* and *R-I.* Solid lines show *R*_1_*, l*_1_and *R*_1_*- l*_1_and dashed lines show *R*_2_*, l*_2_and *R*_2_*- l*_2_. The divergence of *R*_1_and *l*_1_is accompanied with the emergence of the narrow waist in the network.

**Fig. S4.**
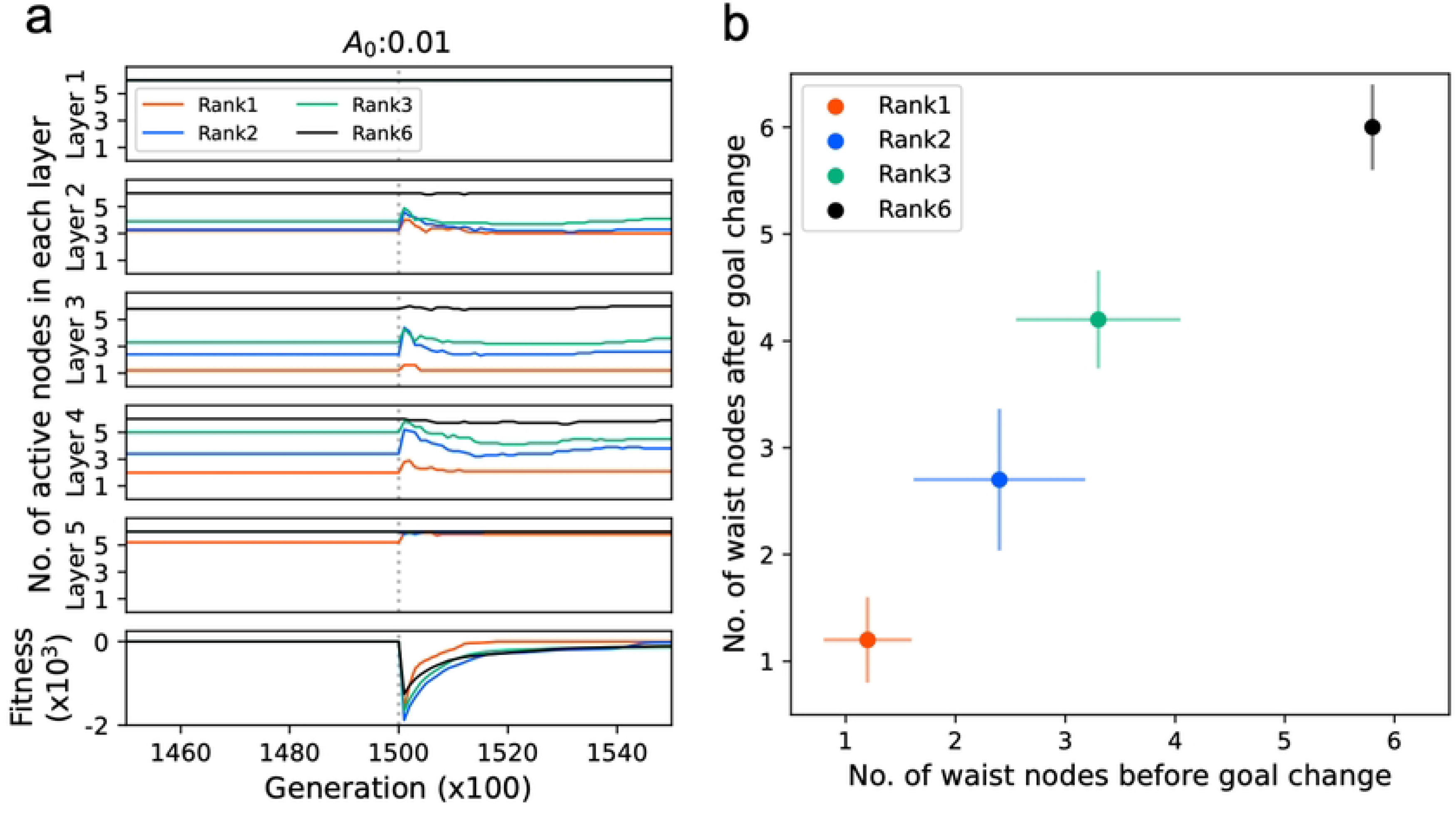
Bow-tie architecture is maintained after changing the goal matrix. (**a)** Evolutionary trajectories of the network from the initial norm *A*_0_ = 0.01. Goal matrix elements are randomized under the conditions of rank1 (red), rank2 (blue), rank3 (green), and rank6 (black). Trajectories are averaged among independent simulation runs (n=10). Y-axis: Number of nodes in each layer. The initial norm *A*_0_ is 0.01. The goal matrix is altered at 150,000 generation (dashed line). (**b)** Comparison between the first and second adapted networks. X-axis: waist width of the network that adapted to the first goal matrix. Y-axis: waist width of the network that adapted to the second goal matrix. Each dot is the average among 10 runs, and the error bars are the std.

**Figure S5.**
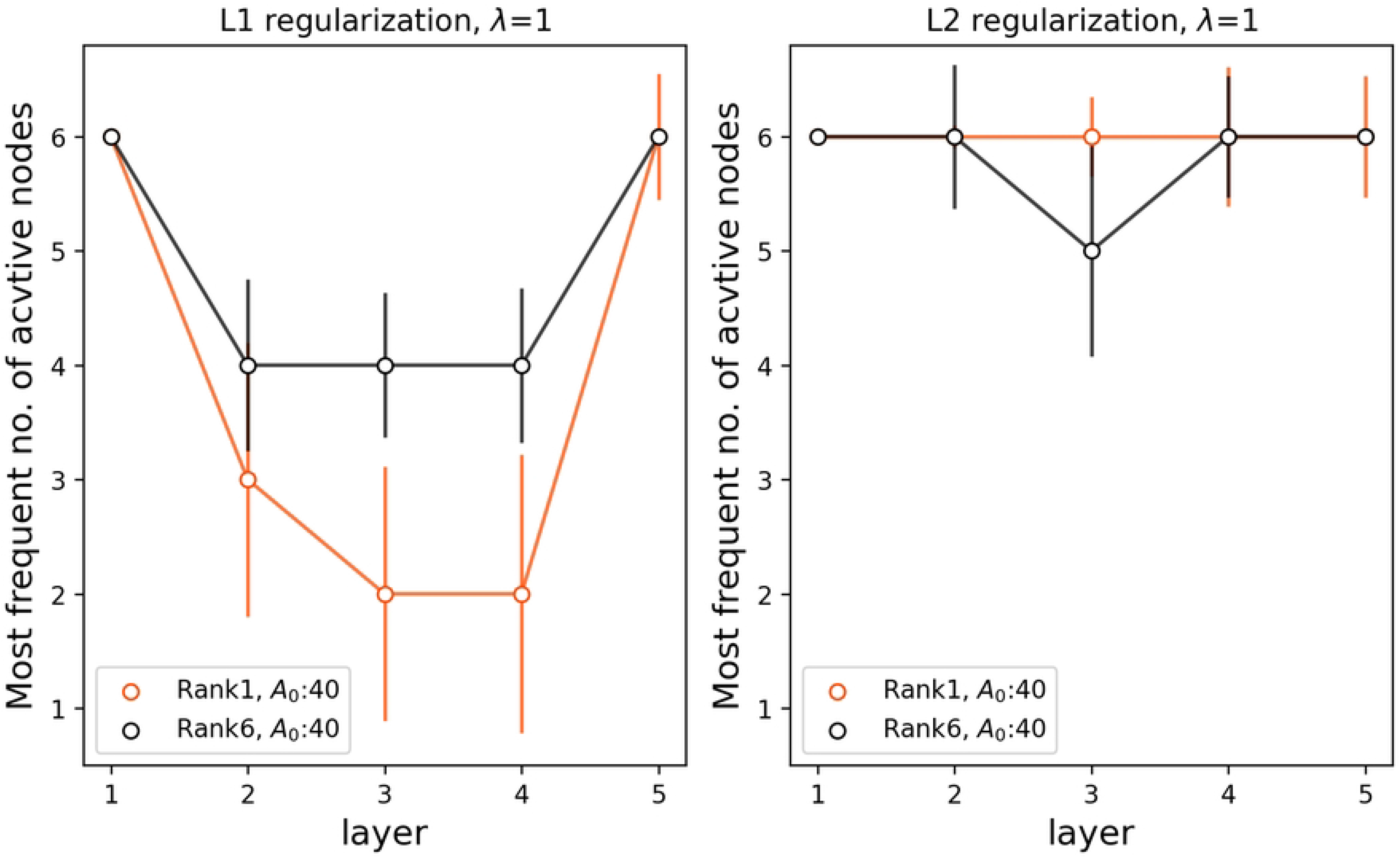
Evolutionary simulation for a bow-tie architecture including the cost for maintaining link intensities. The Y-axis shows the mode among 100 runs of the number of active nodes in the most-adapted network. The number of runs is [rank1: 100; rank6: 100]. Goal matrix elements are randomized under the conditions of rank1(red) and rank6 (black). Initial link intensities are set to *A*_0_: 0.01 (left) and *A*_0_: 40 (right). The error bars represent the std.

## Notes

### Competing Interest Statement

The authors have declared no competing interest.

## References

1. Kitano H. Biological robustness. Nat Rev Genet. 2004;5(11):826–37.

2. Friedlander T, Mayo AE, Tlusty T, Alon U. Evolution of bow-tie architectures in biology. PLoS Comput Biol. 2015;11(3):e1004055.

3. Tieri P, Grignolio A, Zaikin A, Mishto M, Remondini D, Castellani GC, Franceschi C. Network, degeneracy and bow tie. Integrating paradigms and architectures to grasp the complexity of the immune system. Theor Biol Med Model. 2010;7:32.

4. Csete M, Doyle J. Bow ties, metabolism and disease. Trends Biotechnol. 2004 Sep;22(9):446–50.

5. Zhao J, Yu H, Luo JH, Cao ZW, Li YX. Hierarchical modularity of nested bow-ties in metabolic networks. BMC Bioinformatics. 2006;7:386.

6. Stern DL, Orgogozo V. Is genetic evolution predictable? Science. 2009;323(5915):746-51.

7. Mann RS, Carroll SB. Molecular mechanisms of selector gene function and evolution. Curr Opin Genet Dev. 2002;12(5):592–600.

8. Ma HW, Zeng AP. The connectivity structure, giant strong component and centrality of metabolic networks. Bioinformatics. 2003;19(11):1423–30.

9. Ma H, Sorokin A, Mazein A, Selkov A, Selkov E, Demin O, Goryanin I. The Edinburgh human metabolic network reconstruction and its functional analysis. Mol Syst Biol. 2007;3:135.

10. Natarajan M, Lin KM, Hsueh RC, Sternweis PC, Ranganathan R. A global analysis of cross-talk in a mammalian cellular signalling network. Nat Cell Biol. 2006;8(6):571–80.

11. Behar M, Hoffmann A. Understanding the temporal codes of intra-cellular signals. Curr Opin Genet Dev. 2010;20(6):684–93.

12. Jordan JD, Landau EM, Iyengar R. Signaling networks: the origins of cellular multitasking. Cell. 2000;103(2):193–200.

13. Abd-Rabbo D, Michnick SW. Delineating functional principles of the bow tie structure of a kinase-phosphatase network in the budding yeast. BMC Syst Biol. 2017;11(1):38.

14. Supper J, Spangenberg L, Planatscher H, Dräger A, Schröder A, Zell A. BowTieBuilder: modeling signal transduction pathways. BMC Syst Biol. 2009;3:67.

15. Beutler B. Inferences, questions and possibilities in Toll-like receptor signalling. Nature. 2004;430(6996):257-63.

16. Kitano H, Oda K. Robustness trade-offs and host-microbial symbiosis in the immune system. Mol Syst Biol. 2006;2:2006.0022.

17. Polouliakh N, Nock R, Nielsen F, Kitano H. G-protein coupled receptor signaling architecture of mammalian immune cells. PLoS One. 2009;4(1):e4189.

18. Oda K, Matsuoka Y, Funahashi A, Kitano H. A comprehensive pathway map of epidermal growth factor receptor signaling. Mol Syst Biol. 2005;1:2005.0010.

19. Barradas Ana. Evolution of the GPCR signalling system by duplication and its association with disease[Doctoral dissertation]. University of Manchester; 2019.

20. Akhshabi S, Dovrolis C. The evolution of layered protocol stacks leads to an hourglass-shaped architecture. SIGCOMM-Computer Communication Review. 2011;41:206.

21. Broder A, Kumar R, Maghoul F, Raghavan P, Rajagopalan S, et al. Graph structure in the web. Computer networks. 2000;33:309–320.

22. Yan J, Deforet M, Boyle KE, Rahman R, Liang R, Okegbe C, Dietrich LEP, Qiu W, Xavier JB. Bow-tie signaling in c-di-GMP: Machine learning in a simple biochemical network. PLoS Comput Biol. 2017;13(8):e1005677.

23. Wang D, Jin S, Zou X. Crosstalk between pathways enhances the controllability of signalling networks. IET Syst Biol. 2016;10(1):2–9.

24. Ni B, Ghosh B, Paldy FS, Colin R, Heimerl T, Sourjik V. Evolutionary Remodeling of Bacterial Motility Checkpoint Control. Cell Rep. 2017;18(4):866–877.

25. Wells JA. Additivity of mutational effects in proteins. Biochemistry. 1990;29(37):8509–17.

26. Maerkl SJ, Quake SR. A systems approach to measuring the binding energy landscapes of transcription factors. Science. 2007;315(5809):233-7.

27. Friedlander T, Mayo AE, Tlusty T, Alon U. Mutation rules and the evolution of sparseness and modularity in biological systems. PLoS One. 2013;8(8):e70444.

28. Tong AH, Evangelista M, Parsons AB, Xu H, Bader GD, Pagé N, et al. Systematic genetic analysis with ordered arrays of yeast deletion mutants. Science. 2001;294(5550):2364-8.

29. Chauhan P, Shukla D, Chattopadhyay D, Saha B. Redundant and regulatory roles for Toll-like receptors in Leishmania infection. Clin Exp Immunol. 2017;190(2):167–186.

30. Katoch S, Chauhan SS, Kumar V. A review on genetic algorithm: past, present, and future. Multimed Tools Appl. 2021;80(5):8091–8126.

31. Huang Y, Zheng Y, Su Z, Gu X. Differences in duplication age distributions between human GPCRs and their downstream genes from a network prospective. BMC Genomics. 2009;10 Suppl 1(Suppl 1):S14.

32. Vallabhajosyula RR, Chakravarti D, Lutfeali S, Ray A, Raval A. Identifying hubs in protein interaction networks. PLoS One. 2009;4(4):e5344.

33. Bauer R, Kaiser M. Nonlinear growth: an origin of hub organization in complex networks. R Soc Open Sci. 2017;4(3):160691.

34. Hwang JI, Moon MJ, Park S, Kim DK, Cho EB, Ha N, et al. Expansion of secretin-like G protein-coupled receptors and their peptide ligands via local duplications before and after two rounds of whole-genome duplication. Mol Biol Evol. 2013;30(5):1119–30.

35. He X, Zhang J. Why do hubs tend to be essential in protein networks? PLoS Genet. 2006;2(6):e88.

36. Jeong H, Mason SP, Barabási AL, Oltvai ZN. Lethality and centrality in protein networks. Nature. 2001;411(6833):41-2.

37. By Irina Rish, Genady Grabarnik. Sparse modeling: theory, algorithms, and applications. Chapman and Hall/CRC; 2014.

